# MIGRENE: The toolbox for microbial and individualized GEMs, reactobiome and community network modelling

**DOI:** 10.1101/2023.09.01.555866

**Authors:** Gholamreza Bidkhori, Saeed Shoaie

## Abstract

**Motivation:** Understanding microbial metabolism is crucial for evaluating shifts in human host-microbiome interactions during health and disease. However, the primary hurdle in the realm of constraint-based modeling and genome-scale metabolic models (GEMs) pertaining to host-microbiome interactions lies in the efficient utilization of metagenomic data for constructing GEMs that encompass unexplored and uncharacterized genomes. Challenges persist in effectively employing metagenomic data to address individualized microbial metabolism to investigate host-microbiome interactions.

**Result:** To tackle this issue, we have created a computational framework designed for personalized microbiome metabolism. This framework takes into account factors such as microbiome composition, metagenomic species profiles and microbial gene catalogues. Subsequently, it generates GEMs at microbial level and individualized microbiome metabolism, including reaction richness, reaction abundance, reactobiome, individualized reaction set enrichment (iRSE), and community models.

**Availability and implementation:** The MIGRENE toolbox is freely available at our GitHub repository https://github.com/sysbiomelab/MIGRENE. Comprehensive tutorials on the GitHub were provided for each step of the workflow.

## 1. Introduction

Constraint-based modeling, utilizing genome-scale Metabolic Models (GEMs), offers a mechanistic insight into microbial metabolism and genotype-phenotype correlations within host-microbiome interactions (O’Brien et al. 2015). GEMs have been employed for rudimentary microbial community modeling within the gut, showcasing their capacity to discern the pivotal metabolic contributions of bacteria in an ecosystem to the host’s metabolism. Previously, multiple GEMs of human gut bacteria have been formulated, predominantly relying on available whole-genome sequencing data. These models have been instrumental in exploring bacterial growth rates under varying nutrient availability on a community level (Tramontano et al. 2018, Magnusdottir et al. 2017). Nevertheless, a significant challenge endures—specifically, the capability to directly harness metagenomic data for the reconstruction of GEMs that encompass uncultured and not yet characterized genomes, subsequently employing these models for personalized microbiome investigation. Recently, Francisco Zorrilla et al. introduced metaGEM for GEM reconstruction from metagenomes (Zorrilla et al. 2021). Despite this advancement, the hurdle remains in effectively applying metagenomics data for both individual-level microbial metabolism and community-level metabolic modeling.

Employing the latest development of integrated non-redundant microbial gene catalogues and metagenome binning to acquire high quality assembled genomes represents a comprehensive approach to obtaining high-quality GEMs directly from metagenomic data. Here, we present the toolbox for MIcrobial and individualized <u>G</u>EMs, <u>RE</u>actobiome and community NEtwork modelling (MIGRENE), which enables the generation of species and community-level models to be applied to personalized microbiome studies.

## 2. Method

The MIGRENE toolbox effectively utilizes any non-redundant microbial gene catalogue and metagenome species (MSP) to create species-specific Genome-Scale Metabolic Models (GEMs). The initial phase of the workflow involves generating a generalized microbiome GEM through the integration of the non-redundant microbial gene catalogue and a metabolic model (Figure 1a). The reaction profile and reaction score for each metagenome species are established by mapping to the reference model and utilizing taxonomical information (Figure 1b). Subsequently, species-specific GEMs are reconstructed based on the reaction score and bibliomic data (Figure 1c). The analytical component of the toolbox operates using these GEM models, and it’s also possible to incorporate other microbial GEMs into the metagenomics data to enable “individualized metabolic microbiome analysis.” This process generates five distinct outputs at an individualized level: reactobiome, reaction abundance, reaction richness, community modeling, and individualized reaction set enrichment (iRSE) (Figure 1d).

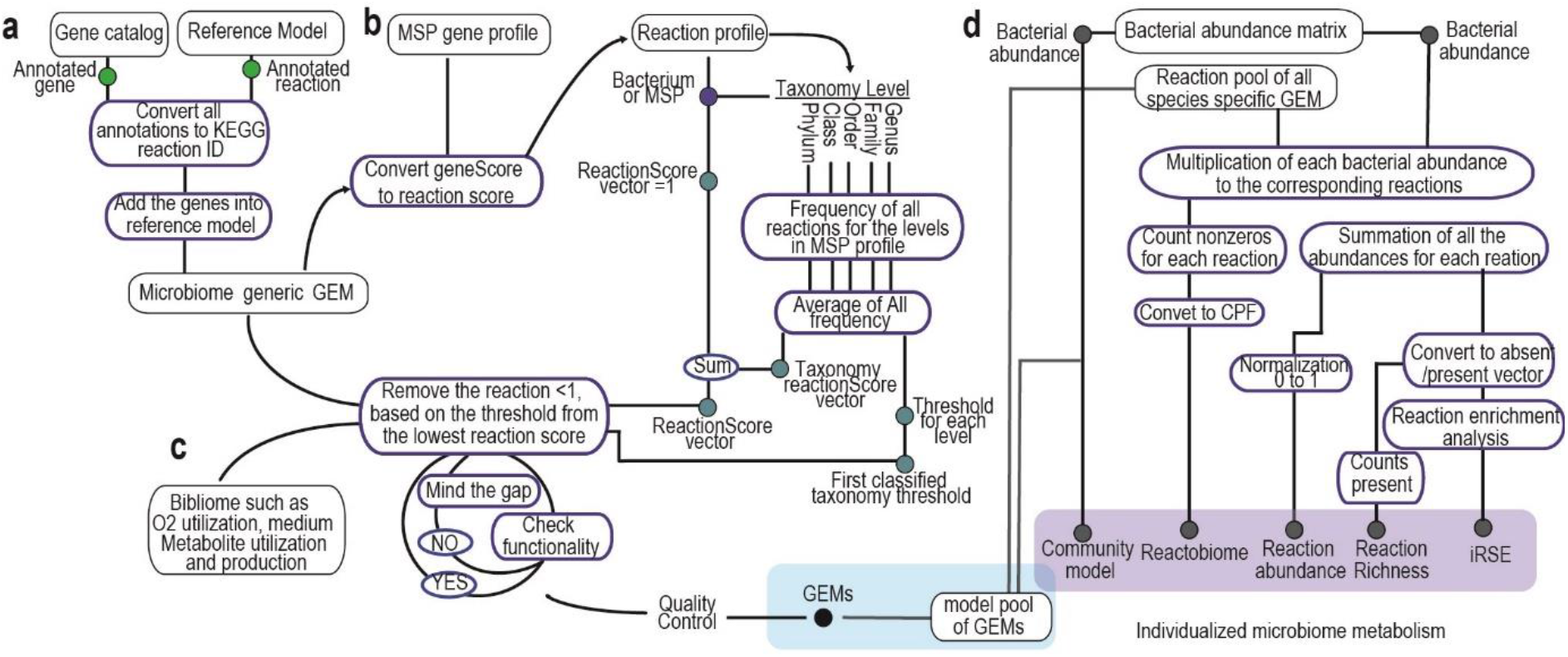
MIGRENE toolbox workflow. **a**, integration of bacterial gene catalog into metabolic model and microbiome reference GEM generation. **b-c**, calculation of reaction score and species-specific model)generation. **d**, generating individualized microbiome metabolism.

### 2.1 integration of bacterial gene catalog into metabolic model and microbiome reference GEM generation

First, the bacterial gene catalogue undergoes restructuring and verification using the “checkCatalog.m” function. The “convertCatalogAnnotation.m” function is then employed to translate KO annotations from the catalogue into KEGG reaction IDs. If the mapping ID file is not specified as an input argument, the function automatically fetches the required conversion information from the KEGG API and stores it in the “data” directory within the MIGRENE Toolbox location.

Next, a comprehensive annotated metabolic model is generated by leveraging the resources of the generic metabolic model (such as ModelSEED and KBase). Both the generic metabolic model and the gene catalogue serve as inputs for the “microbiomeGEMgeneration.m” function, facilitating the creation of a reference genome-scale metabolic model (GEM). The association between genes, proteins, and reactions (Gene-Protein-Reaction or GPR) is established by integrating the catalog genes into the metabolic model using KEGG reaction IDs. In cases where the GEM’s annotation file is not provided, the function autonomously identifies annotations such as KO, reaction ID, or EC within the model and converts them into reaction IDs based on the most recent version of KEGG IDs. Additional fields, namely “grRules,” “genes,” “rules,” “geneNames,” and “rxnGeneMat,” are introduced into the GEM to ensure compatibility with both COBRA and RAVEN GEMs. Consequently, the resulting models can be seamlessly employed with the functions offered by both toolboxes.

### 2.2 Calculation of reaction score and taxonomy threshold

The annotated reference GEM was used to convert the MSP gene profile to reaction profiles. The reaction states (absent/present) for each MSP were generated based on the presence of MSP genes in the corresponding gene rule vector of the reaction in the reference GEM. A matrix of reaction frequency was also calculated using the average of frequency of each reaction in the taxonomy levels (*i*.*e*. genus, family, order, class, phylum) of the corresponding MSP. Reaction scores for each MSP were calculated by adding the reaction state and reaction frequency matrices entries together, in result the reaction score profiles for the MSPs were made. A threshold (the lowest non-zero frequency) was determined for each taxonomy level for filtering the reactions while generating the species-specific GEMs (pseudocode is available in Supplementary note). MetagenomeToReactions.m function creates a reaction and the reaction states for all the MSPs and taxonomy information are collected using GenerateMSPInformation.m. MetaGenomicsReactionScore.m calculates reaction scores for each MSP (bacterial species)

### 2.3 species-specific genome-scale metabolic model generation

contextSpecificModelGenertion.m and then contextSpecificModelTune.m are used to generate the GEM models. Instead of regular gap filling, the former function pruned the constrained reference GEM from the lowest reaction score and minded the gaps to keep the model functional based on biomass objective function and the threshold. So, we started gap filling from the reactions with the highest frequency in the closest taxonomy level of the species. The latter function tuned the models by pruning the exchange and transport reactions, dead end metabolites, adding compartments, model description and correcting KEGG metabolite ID and metabolite formula. It also calculated the important information for gap filling percentage at different levels: i.e. taxonomy proximity based, further taxonomy level, not annotated (without gene-protein-reaction relation) and the total gap filling percentage. The growth rate for each model were tested for the diets using constraint-based modelling. Additionally, the structure, connectivity and carbon balance for the generated GEMS are provided in the toolbox.

### 2.4 generation of individualized metabolic microbiome: Reaction richness, reaction abundance and reactobiome, iRSE and community models

We utilize a reaction pool from GEMs and the abundance of Metagenome Species (MSP) to create personalized microbiome metabolism. Reaction richness profiles, representing gut microbiome compositions, are computed using the RxnRichnessGenerator.m function. Each MSP’s abundance contributes to the reactions of the sample, forming a matrix (n×m; n: reaction pool size, m: MSP count) for each sample. This matrix becomes a binary vector, indicating the presence/absence of reactions in samples, capturing reactions present in at least one MSP (supplementary figure). The ReactionAbundanceGenerator.m sums reaction abundances to yield relative reaction abundance profiles. Reactobiome represents normalized reactions per 500 bacteria (CPF) in the gut for each individual, calculated using CPFGenerator.m. Individualized reaction set enrichment (iRSE) is generated by pRSEGenerator.m, showing overrepresentation of 153 KEGG metabolic pathways (provided in the toolbox) in samples based on a Hypergeometric test and a binary vector.

Community models are created for each sample using MakeCommunity.m. Bacterial species names are added as prefixes to reaction names in GEMs, segregating species-specific GEMs in the combined S matrix. Compartments [lu] (intestinal lumen) and [fe] (secreted bacterial and food-derived metabolites) are included, connected by exchange reactions. [lu] represents the gut microbiome with bacterial GEMs and exchange reactions for metabolites. Community biomass objective function accounts for bacterial biomass, with bacterial abundance as stoichiometric coefficients.

## 3. Results and discussion

We have developed the MIGRENE Toolbox, a comprehensive platform designed to seamlessly integrate microbial gene catalogues, metagenomic species data and general metabolic models. This integration empowers the generation of Genome-Scale Metabolic Models (GEMs) and community-level models, facilitating their application in individualized microbiome studies.

The utility of this toolbox has been prominently demonstrated in a variety of recent studies. One remarkable application involved the exploration of the mechanistic implications of the gut microbiome in three distinct metabolic disorders: obesity, type 2 diabetes, and atherosclerosis. Through the utilization of the MIGRENE Toolbox, an in-depth investigation revealed noteworthy patterns. Specifically, a shared phenomenon emerged across these disorders – a heightened propensity for glutamate consumption, coupled with the concomitant production of ammonia, arginine, and proline by gut bacteria (Proffitt et al. 2022). Another instance highlighting the toolbox’s prowess emerged from the work of Ezzamouri et al. (Ezzamouri et al. 2023). This study successfully established a compelling association between the human gut microbiome and type 2 diabetes mellitus. The researchers further delved into the intricate changes in bacterial metabolism following metformin treatment, a key aspect in diabetes management. Furthermore, the MIGRENE Toolbox was used in illuminating the microbial landscape in Parkinson’s disease (PD). Researchers unveiled a heightened microbial capacity for mucin and host glycan degradation and folate deficiency within the context of PD.

## Funding

This study was supported by Engineering and Physical Sciences Research Council (EPSRC), EP/S001301/1, Biotechnology Biological Sciences Research Council (BBSRC) BB/S016899/1, Science for Life Laboratory, the Knut and Alice Wallenberg Foundation, the Erling Persson Foundation and King’s College London.

## Conflict of interest

The authors declare no competing interests.

## Notes

https://github.com/sysbiomelab/MIGRENE

### Competing Interest Statement

The authors have declared no competing interest.

