## SUPPLEMENTARY DATA for "MIGRENE: The toolbox for microbial and individualized GEMs, reactobiome and community network modelling"

### Supplementary note

#### Calculation of reaction score (pseudocode)

```
INPUT referenceGEM
INPUT mspGene (binary n×m matrix, n: number of genes; m: number of MSPs)
INPUT taxonomy (m×r table, m:number of MSPs; r: number of taxonomy level i.e. from genus to phylum,
[m,r]: taxonomy name)
reactionState= zeros (length(referenceGEM.rules), m)
FOR h = 1 to m
        vect= gene names (mspGene(:,h)==1)
        FOR i=1 to length(vect)
                index=find (vect_i in referenceGEM.rules)
                IF NOT empty (index) THEN reactionState(index,i)=1 ENDIF
        ENDFOR
ENDFOR
% convert reaction state to reaction score
FOR i = 1 to m
        FOR j= 1 to r
                IF taxonomy (i,j) != "unclassified" THEN
                        index=find (taxonomy (i,j) in taxonomy (:,j))
                        state= reactionState (:,index)
                        FOR k= 1 to length(referenceGEM.rules)
                                ScoreMatrix(k,j) = sum(state(k,:)) /r;
                         ENDFOR
                ELSE
                        ScoreMatrix(:,j)=0
                ENDIF
        ENDFOR
        FOR k= 1 to length(referenceGEM.rules)
                ScoreMatrix(k,(r+1)) = sum(ScoreMatrix (k,:)) /r;
        ENDFOR
        reactionScore(:,i)= ScoreMatrix(k,(r+1))
ENDFOR
FOR i = 1 to length(referenceGEM.rules)
        IF empty (referenceGEM.rules(i)) THEN
                reactionState(i,:)=-1; reactionScore(i,:)=-1
        ENDIF
ENDFOR
% Define threshold
FOR j= 1 to r
threshold (1,j)= MIN ScoreMatrix(1,j)>0
ENDFOR

RETURN  threshold
RETURN  reactionScore
RETURN  reactionState
```

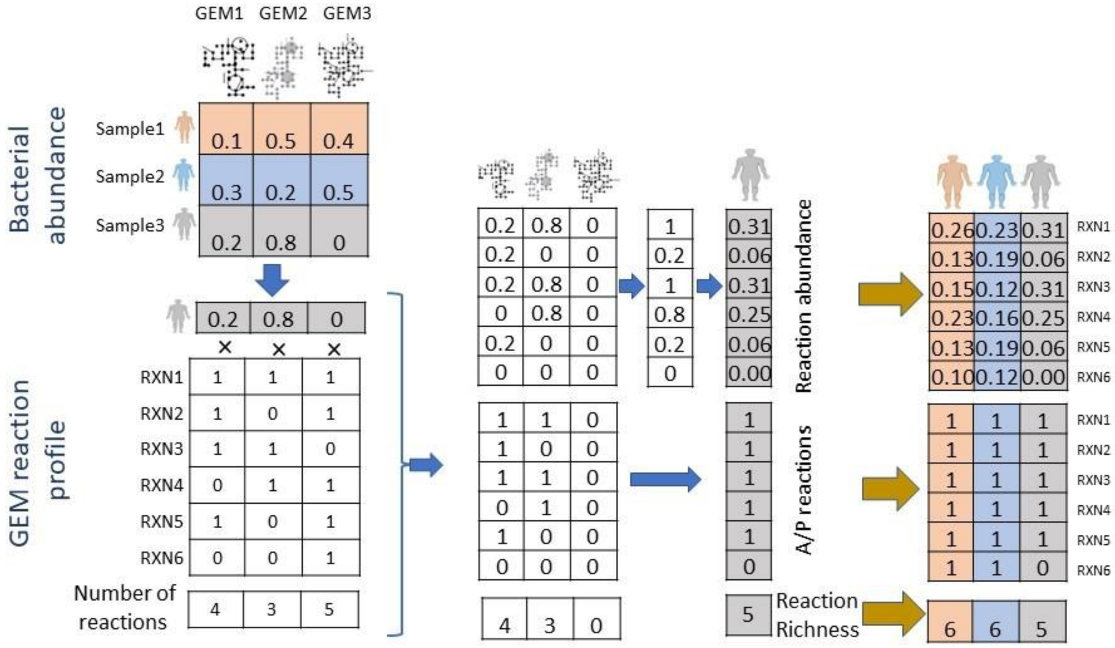


**Supplementary figure.** Calculation of reaction abundance and richness for 3 individuals using 3 bacterial abundance and the corresponding GEMs. rxn: reaction; A/P: absent/present.
